# Extracting the GEMs: Genotype, Environment and Microbiome interactions shaping host phenotypes

**DOI:** 10.1101/863399

**Authors:** Ben O. Oyserman, Viviane Cordovez, Stalin W. Sarango Flores, Harm Nijveen, Marnix H. Medema, Jos M. Raaijmakers

**Affiliations:** Department of Microbial Ecology, Netherlands Institute of Ecology, 6708 PB Wageningen, The Netherlands; Bioinformatics Group, Wageningen University & Research, 6708 PB Wageningen, The Netherlands; Institute of Biology, Leiden University, 2333 BE Leiden, The Netherlands; Colegio de Ciencias Biológicas y Ambientales, Universidad San Francisco de Quito, Quito, Ecuador

## Abstract

One of the fundamental tenets of biology is that the phenotype of an organism (*Y*) is determined by its genotype (*G*), the environment (*E*) and their interaction (*GE*). Quantitative phenotypes can then be modeled as *Y*=*G*+*E*+*GE*+*e*, where *e* is the biological variance. This simple and tractable model has long served as the basis for studies investigating the heritability of traits and decomposing the variability in fitness. Increasingly, the importance of microbe interactions on organismal phenotypes is being recognized, but it is currently unclear what the relative contribution of microbiomes to a given host phenotype is and how this translates into the traditional GE model. Here we address this fundamental question and propose an expansion of the original model, referred to as GEM, which explicitly incorporates the contribution of the microbiome (*M*) to the host phenotype, while maintaining the simplicity and tractability of the original GE model. We show that by keeping host, environment and microbiome as separate but interacting variables, the GEM model can capture the nuanced ecological interactions between these variables. Finally, we demonstrate with an *in vitro* experiment how the GEM model can be used to statistically disentangle the relative contributions of each component on specific host phenotypes.

## The genetic basis of ecological interactions

Leveraging the beneficial interactions between plant hosts and their microbiomes represents a new direction in sustainable crop production. In particular, the emergence of *m*icrobiome-*a*ssociated *p*henotypes (MAPs) (Oyserman *et al.*, 2018), such as growth promotion and disease suppression, is expected to reduce our dependency on energy-intensive and environmentally disturbing management practices. This may either be achieved through the addition of probiotics and prebiotics, or through breeding programs targeting MAPs to develop a next generation of ‘microbiome-activated’ or ‘microbe-assisted’ crop production systems (Busby *et al.*, 2017; Oyserman *et al.*, 2018). Hence, a major challenge is to identify the genotypic underpinning of emergent MAPs and understanding the pivotal role of the environment. To date, however, the relative contribution of microbiomes to a given host phenotype is not known for most host phenotypes. The interaction between genotype (G) and environment (E) has long been recognized as an important factor both in evolutionary biology (Via & Lande, 1985; Anderson *et al.*, 2013) and breeding programs (Allard & Bradshaw, 1964). While a significant body of literature exists on quantitative investigations of GE interactions (El-Soda *et al.*, 2014), the bulk of this work has focused on abiotic parameters and has largely overlooked the microbiome. Nevertheless, the interactions between hosts, microbiomes and their environments are coming into increasing focus and scrutiny (Dal Grande *et al.*, 2018; Wallace *et al.*, 2018; Beilsmith *et al.*, 2019; Bonito *et al.*, 2019).

One current opinion is that rather than viewing host plants and animals as individuals, they should be viewed together with their microbiomes as single cohesive unit of selection termed a ‘holobiont’ with a ‘hologenome’(Bordenstein & Theis, 2015; Moran & Sloan, 2015; Douglas & Werren, 2016). Under this view, the microbiome (M) could be integrated into the G term of the GE model of host phenotypes. However, others have pointed out that treating hosts and their microbiomes as a single unit does not capture the broad range of interactions and fidelity between host and microbe (Douglas & Werren, 2016). Another popular opinion is that, as the environment is classically defined to include “physical, chemical, and biotic factors (such as climate, soil, and living things) that act upon an organism” (‘Environment’, 2019), M should be integrated into the E term of the GE model. However, an important distinction exists between E and M components; M is dynamic (i.e., have many interdependencies and may adapt or evolve through time), while E is driven through external processes. Here, we address these two viewpoints and propose that it is useful to introduce microbiomes and MAPs as a discrete unit within the GE model. In doing so, we put forth an updated GEM model that explicitly incorporates the microbiome (M) and its respective interactions with the genotype (G) and environment (E). Using these mathematical representations, we conceptually emphasize interesting cases that emerge from this framework (Figure 1). Finally, we present a simple ‘one-microbe-at-a-time’ experiment to highlight key features and challenges of unearthing GEM interactions, and to statistically disentangle the relative contributions of each of the GEM model components (Figure 2).

**Figure 1.**
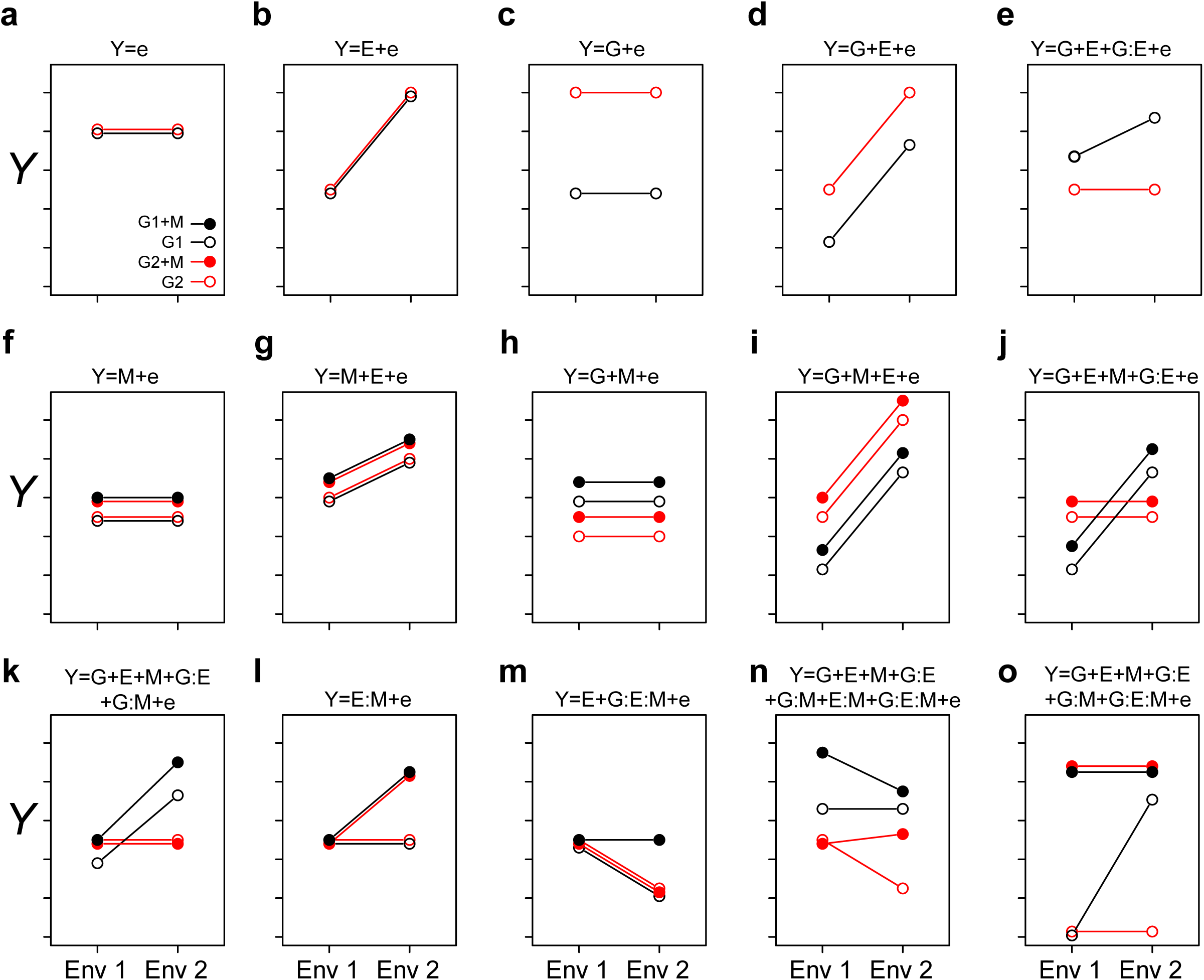
Conceptualizing the GEM model: Here we graphically explore how the interactions between genotypes, environment and microbiome may impact a host phenotype (Y). The two genotypes are indicated by G1 and G2, and the presence of a microbiome is indicated by solid circles (as shown in panel a). The different environments are indicated as Env 1 and Env 2 on the X-axis. In each case (panels a-o), the corresponding equation is depicted over the figure itself. In cases when we treat the microbiome as a phenotype of the host, the relative abundance of a particular taxon, or other features of a microbiome, may be considered as the sum of G and E interactions (panels a-e). In simple cases, the relative abundance is independent of genotype (panel b) or environment (panel c). More likely, both genotype and environment, and their interactions will contribute to relative abundance/function (panels d and e respectively). Panels a-e are special cases of the GEM model, indicating situations in which the microbiome does not contribute to a particular host phenotype. Building complexity, each of G, E and M may contribute to host phenotypes individually or in combination, but without interaction (panels a-d and f-i). Finally, the highest level of complexity occurs once interactions between G, E and M occur (panels e, j-o). A salient feature of this representation is that when no interaction between variables exists, the slope is equal between treatments. This model may also provide practical insights, such as identifying optimal prebiotics which may be expected to have a broad host range (no G interaction) and be conditionally neutral (panel l). Additionally, this model may serve to characterize complex interactions, such as conditional symbiosis where a host fitness is reduced to zero without a microbiome (taxon or function) in a particular environment (panel o).

**Figure 2.**
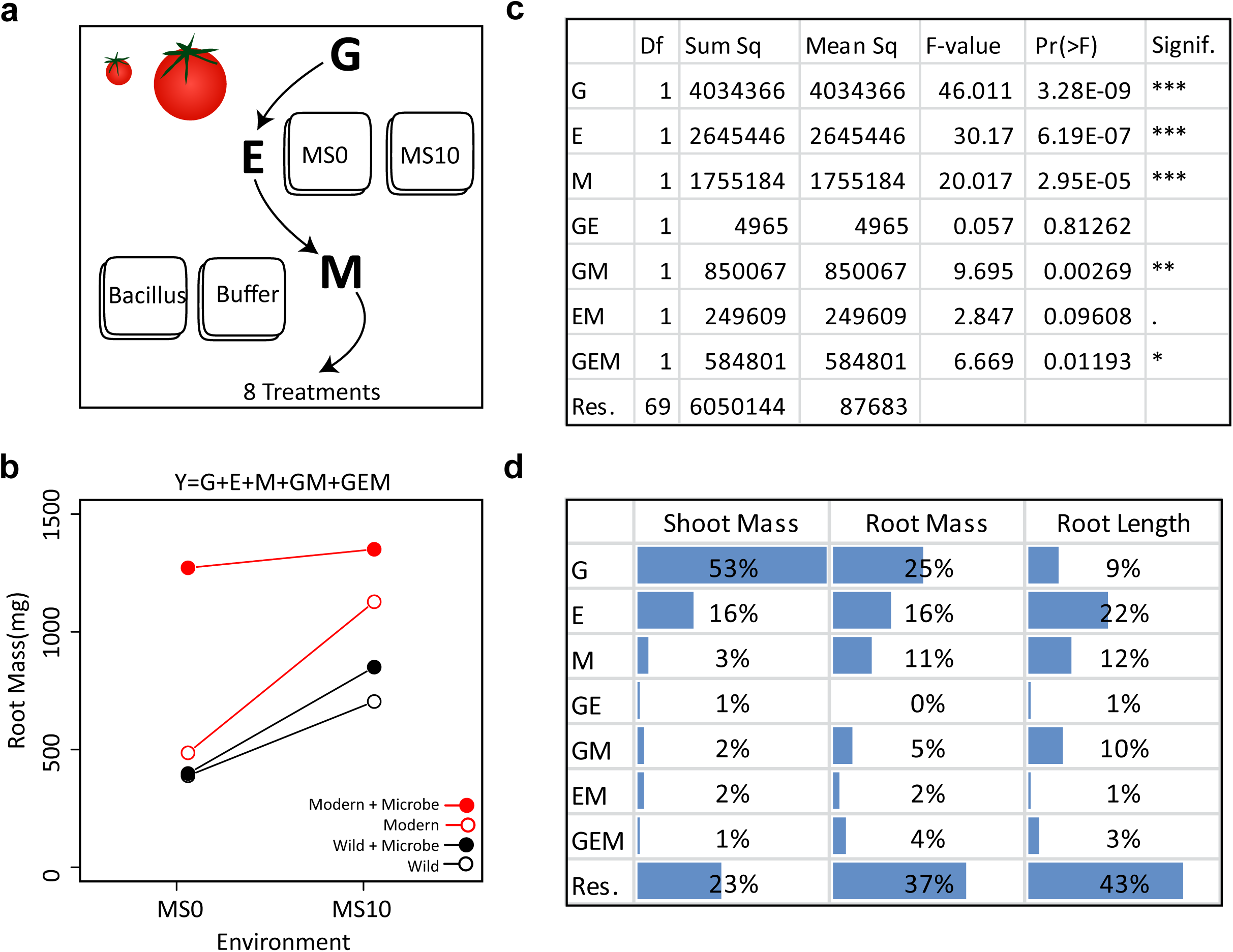
Extracting the GEMs from the simplified GEM experiment: (Panel a) In this in vitro experiment, the contribution of G, E, M and their interactions were investigated in a fully factorial design. (Panel b) In total, two tomato genotypes, two environments and one microbe treatment were investigated. Various plant phenotypes were measured, but for clarity, only the average dry root mass of each treatment are visualized here. (Panel c) The GEM model shows that G, E, M, GM and GEM all contribute significantly to root mass. The ANOVA table displays the reported Df (Degrees of freedom), Sum sq (Sum-of-squares), Mean sq (Mean some-of-squares), the F-value (the test statistic of an ANOVA), Pr(>F) (the p-value), and Signif. (a visual indication of the level of significance). (Panel d) Here we present the ANOVA outcome showing the percent of the total sum of squares for dry shoot mass, dry root mass and root length. For shoot mass, plant genotype explained the greatest portion of variance. In contrast, both E and M explained a greater amount of variation than plant genotype for root length. Importantly, for each of the three plant phenotypic parameters measured, GM explained a greater amount of variation than GE.

## The microbiome as a phenotype or microbiome-associated phenotypes?

The relationship between the host and its microbiome may be generally defined and viewed in two ways. Firstly, microbiome community structure may be considered a phenotype of the host (*Y*), henceforth ‘microbiome as a phenotype’(Belheouane *et al.*, 2017; Rothschild *et al.*, 2018; Walters *et al.*, 2018). Under this view, taxonomic/functional features of the microbiome, are treated as the phenotype of the host (*Y*). In this manner, *Y* (e.g. the abundance of a taxon or functional gene) may be represented based on the contribution and interaction between the genotype (*G*), the environment (*E*) and the remaining variance (*e*) (Equation 1).

Secondly, a microbiome may be quantified by their impact on the host phenotypes (Kopac & Klassen, 2016; Oyserman *et al.*, 2018). In this view, MAPs such as plant growth promotion or plant health are treated as the phenotype (*Y*) (Zeevi *et al.*, 2019). Here, we suggest explicitly expanding the environmental parameter of the traditional GE model (Equation 1), such that host genotype (*G*), environmental factors (*E*) and microbiome structure and function (*M*) and their interactions all contribute to the observed host phenotype (Equation 2). Thus, measurements of the microbiome structure and function are used in conjunction with genotypic and environmental data to explain a MAP, an emergent phenotype of the host-microbe interaction. Additional components may be added to the GEM model to accommodate additional complexity. For example, M may be split into *i* components, where M*i* represents the *i*^th^ taxonomical or functional feature. In this way, the GEM model is amenable for investigating the role of microbe-microbe interactions within natural or synthetic communities, the interactions between multiple environmental factors, or any complex arrangements (see supplemental materials for discussion on an expanded GEM model). In Figure 1, we exhibit some basic features of the GEM model.

## Extracting the GEMs

To demonstrate how the GEM model may be used to disentangle the relative influence of various factors on a particular host phenotype, we investigated GEM interactions in a simplified *in vitro* assay with one bacterial strain (*Bacillus* sp., accession number MN512243) interacting with two plant genotypes, a modern domesticated tomato cultivar (*Solanum lycopersicum* var moneymaker) and a wild tomato relative (*Solanum pimpinellifolium*) under two environmental conditions. In this model system, all genotype, environmental, microbial parameters are controlled and therefore can be systematically explored in a fully factorial design (details are in the supplemental material). For each tomato genotype, seedlings were grown in two environments, i.e. Murashige and Skoog agar medium (MS0) and MS agar medium supplemented with 10 g/L of sucrose (MS10). After germination, the root tips were inoculated with the *Bacillus* strain, which was originally isolated from the wild tomato rhizosphere. Control seedlings were inoculated with buffer only (Figure 2A). The plant phenotypes monitored were root architecture (using WinRhizo™) and root and shoot dry mass (Figure 2B). An ANOVA was done to test the significance of each variable in the GEM model (Figure 2C). Together, the microbiome (M) and all interacting variables (GM, EM and GEM) explained 26% of root dry mass variance, 21% of shoot dry mass variance and 8% of root length total variance. Furthermore, in all cases the interacting parameters, GM, EM, and GEM interactions explained greater variance than GE interactions (Figure 2D).

A clear consensus is forming that microbiomes impact host phenotypes, but its relative contribution to that host phenotype is, in most cases, not known. The GEM model provides a simple, tractable and testable model demonstrating that the interactions of the microbiome and other model terms (GM, EM and GEM) are also essential determinants of host phenotypes. It is important to highlight that, in this case, GM interactions actually explain more variability than canonical GE interactions. Furthermore, the expanded GEM model captures other important features that may otherwise be easily overlooked, such as the genotype-independent interaction between EM. This states that microbe and environment may interact to alter host fitness independent of the genotype. For example, auxin is a plant hormone that promotes growth that is also produced by bacteria. Many bacterial cultures have differential auxin production dependent on their environment (Tsavkelova, 2005); therefore, it is likely that EM interactions can promote auxin production and thus plant growth independent on genotype. In practice, identifying EM may have important implications for synbiotics (mixtures of probiotics and prebiotics). In this manner, the GEM model not only provides a model to disentangle the contribution of G, E and M, but also serves as a powerful tool for conceptualization.

## The GEM model captures complex ecosystem processes

As describe above, genotype, environment and microbiome may influence organismal phenotype directly, but also through their interactions. This dynamic is captured by the various *terms* that make up the GEM model, providing a simple means to conceptualize this otherwise complex system. In its most basic form (Equation 2), the GEM model has 8 terms in total. An example of a term with a single variable is ‘G’, a two variable term would be ‘GM’, and three variable term would be ‘GEM’. While the basic GEM model contains terms related to inter-class interactions (GE, GM, etc.), it lacks terms representative of intra-class interactions (M:M, E:E, etc). By simply adding additional variables to the GEM model, M:M and other ecologically relevant interactions may be introduced as additional terms. The number of terms in a model is dependent on the number of variables (*n*) that can be mathematically represented by Supplemental Equation 1. In addition, the number of terms with *r* variables may be mathematically represented by Supplemental Equation 2, where *n* is the total number of variables, and *r* is the number of variables in the term. From this basis, a model of organismal phenotype which takes into account ecosystem-level processes may be constructed. To this end, we developed a simple Python script to generate a GEM model based on user input for any number of G, E and M variables (https://github.com/Oyserman/GEM).

To model the interactions between multiple microbiome members, such as those found in natural or synthetic communities, in Equation 3, we provide a simple expansion of the basic GEM Equation presented in the main text to add another microbiome variable. The result is a 4 variable (GEM_l_ M_2_) model that includes all *r*-way interactions terms necessary to model the impact of a two member community on any number of plant genotypes or environments. For clarity, Equation 3 is presented with all *r*-way interactions on separate lines. To show the versatility of the GEM model, we provide another expansion in which multiple hosts are interacting in a particular ecosystem (G_1_G_2_EM). In this case, the fitness of one plant genotype (G_1_) is influenced through interactions with a neighboring plant genotype (G_2_) and their associated microbiomes. A prominent example of this in literature are intercropping systems in which nitrogen fixation through legume-microbiome interactions benefit other non-leguminous plants in a nitrogen limited soil ecosystem (Peoples *et al.*, 1995).

## Conclusions

A fundamental tenet of biology is that genotype and environment interact and impact the fitness and phenotype of an organism. The GE model of organismal phenotype has been the cornerstone of modern breeding programs. Part of the power of the GE model is its simplicity and interpretability. However, the important role of host-associated microbiomes has recently come into focus. Here, we investigated how microbiomes (M) fit into the GE model, suggest an explicit expansion to include M, and argue that, because of its dynamic and evolving nature, that M should not be collapsed within E. We use a conceptual figure to show that the updated GEM model captures the diverse possible outcomes of between G, E and M. To support our model, we present an *in vitro* experiment with one microbe demonstrating not only how to use the GEM model, but also showing that GM interactions may explain more variability than GE interactions. Finally, additional examples of expanded GEM models which take into account M:M and G_2_:E:M interactions are presented to demonstrate the ecological versatility of the GEM model. Taken together, we propose that the GEM model provides a simple and interpretable expansion of the GE model. Furthermore, given the important role of the microbiome, any investigations into GE interactions must also account or control for M.

## Conflict of interest

The authors declare that they have no conflict of interest.

## Acknowledgements

We thank Victor Carrion, Azkia Nurfikari and Hannah McDermott for contributing the isolate. The contributions of Ben Oyserman and Jos Raaijmakers were funded in part by the Technology Foundation of the Dutch National Science Foundation (NWO-TTW)

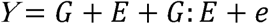

***Equation 1.* The traditional model for GE interactions**: In the canonical model of quantitative phenotypes, the host phenotype (*Y*) is explained by the sum of G, E, their interactions (G:E), and *e* the residual error. This model may be used to calculate the proportion of variance explained by the host genome and the environment on a host associated microbiome community. In other words, the microbiome may be treated as *Y*, the phenotype of the host (e.g. ‘the microbiome as a phenotype’). When E has no contribution to *Y*, only G determines the abundance or function of the microbiome (Figure 1C). On the other side of the spectrum, only E determines to the abundance or function of the microbiome (Figure 1B).

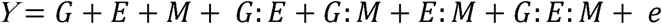

***Equation 2. The new GEM model:*** When a microbiome has a quantitative impact on host phenotype, the traditional GE model may be expanded to incorporate M and all respective interactions (GM, EM, and GEM). Unlike the GE model, which may be used to explain the microbiome, the expanded GEM model may be used to statistically disentangle the contribution of G, E and M and their various interactions to changes in host phenotype. When M has no impact, this variable and those associated with it fall out of the equation giving the GE model. These, and other special cases are conceptually explored further in Figure 2. Thus, this model is capable of capturing the nuanced dynamics of host-microbiome interactions, such as host-microbe interactions that are environment-specific, or otherwise have lower fidelity than strict symbiosis (Douglas & Werren, 2016).

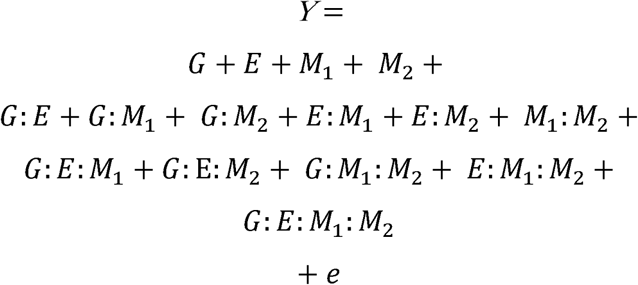

***Equation 3. A GEMM model: The basic GEM model may be expanded to include any number of complex interactions. Here we expand the GEM model to include microbe-microbe interactions. This results in the addition of 1-way, 2-way, 3-way and 4-way interaction terms, which are shown on separate lines for clarity.***

## Notes

https://github.com/Oyserman/GEM

